# Rho2-dependent cell wall remodeling boosts the fungistatic activity of manogepix against *Aspergillus fumigatus*

**DOI:** 10.1101/2025.10.20.683572

**Authors:** Sean Brazil, Isabel Kopp-Fouquet, Samuel Zen Kai Yap, Michael Carty, Johannes Wagener

## Abstract

Fosmanogepix is a new antifungal agent currently undergoing phase 3 clinical trials. Its active moiety, manogepix, inhibits Gwt1 which is an enzyme essential for the assembly and attachment of glycosylphosphatidylinositol anchors to cell wall proteins. Manogepix has a strong fungistatic activity against most human pathogenic fungal species. Here we characterized the activity of manogepix against the major human pathogenic mold *Aspergillus fumigatus*. We show that manogepix susceptibility is linked to the expression of *gwt1*, the gene encoding Gwt1, thereby demonstrating that overexpression of *gwt1* can confer resistance. In agreement with a previous study conducted with a different fungal pathogen, we observed an increase of cell well chitin after manogepix treatment. Using a luciferase-based reporter assay, we show that manogepix activates the cell wall integrity stress response pathway. However, manogepix only occasionally causes cell lysis. Mutants that lack the cell wall stress sensor Wsc1 or the Rho GTPase Rho4, which is important for septum formation, are slightly more susceptible to manogepix. In contrast, mutants that lack the Rho GTPase Rho2 or the cell wall stress sensor MidA, both of which are key for *A. fumigatus* to survive cell wall stress caused by granulocytes, heat and other cell wall perturbing chemical compounds, grow better than wild-type when exposed to manogepix. We show that both mutants, especially Δ*rho2* but also Δ*midA*, fail to upregulate cell wall chitin to wild-type-like levels in response to manogepix. These results implicate a role for the Rho2-dependent stress response in the fungistatic activity of manogepix against *A. fumigatus*.

## INTRODUCTION

*Aspergillus fumigatus* is a mould and airborne opportunistic fungal pathogen. It causes infections in humans, of which invasive aspergillosis is the most feared. Invasive aspergillosis is a severe systemic infection that occurs primarily in immunocompromised patients and is associated with a high mortality rate even when treated. There is currently only a limited number of drugs that are approved for the treatment of infections caused by *Aspergillus* species. These belong to essentially three drug classes: Azoles, which inhibit sterol C14-demethylase CYP51, a key enzyme in the synthesis of the primary fungal sterol ergosterol, resulting in the accumulation of toxic ergosterol precursors (Elsaman et al., 2024; Rybak et al., 2024). Polyenes, which bind to ergosterol having detrimental effects on the membrane integrity, including the formation of fungicidal ion channels (Lewandowska et al., 2021). And, as a second-line medication, echinocandins, which inhibit β-1,3-glucan synthase required for cell wall biogenesis and which exert only a fungistatic activity against *Aspergillus* species (Dichtl et al., 2015). The emergence of resistance to azole antifungals, frequent problems with drug-drug interactions and tissue penetration, and the high toxicity of polyene antifungals highlights the need for new drugs for the treatment of fungal infections.

Fosmanogepix is a new first-in-class antifungal for the use in humans. It shows activity against most human pathogenic fungi, including *Aspergillus* species, and is currently being investigated in phase 3 clinical trials for the treatment of patients with invasive fungal infections (ClinicalTrials.gov, NCT05421858, NCT06925321, 2025). Fosomanogepix is the pro-drug of manogepix (APX001), which inhibits the fungal enzyme Gwt1, located in the endoplasmic reticulum (ER). Gwt1 is required for the assembly and attachment of glycosylphosphatidylinositol (GPI) anchors to proteins destined for the cell surface (Tsukahara et al., 2003; Umemura et al., 2003). Gwt1 catalyzes the palmitoylation of glucosamine-phosphatidylinositol (GlcN-PI) at the second position of the inositol ring using palmitoyl coenzyme A (palmitoyl-CoA) as the substrate (Dai et al., 2024; Tsukahara et al., 2003; Umemura et al., 2003). Inhibition of this enzyme causes an accumulation of immature cell surface proteins in the ER which has fungistatic effects (Mann et al., 2015; McLellan et al., 2012; Miyazaki et al., 2011). Since many GPI-anchored proteins have an integral role in maintaining the structure and organisation of the fungal cell wall (Ibe and Munro, 2021; Samalova et al., 2020), detrimental effects on the cell wall stability are to be expected.

Here we studied the antifungal effects of manogepix against *A. fumigatus*. We show that overexpression of *gwt1* can act as a mechanism to overcome the growth inhibitory effect of manogepix in this mold. Furthermore, we show that manogepix triggers activation of a cell wall salvage response, which results in an increase of cell wall chitin and involves the Rho GTPase Rho2. Surprisingly, this activation contributes to the fungistatic effect of manogepix.

## RESULTS

### Construction and phenotypic characterisation of a conditional *A. fumigatus gwt1* mutant

Manogepix targets the enzyme Gwt1. Homologs of this enzyme have not been studied in *Aspergillus* species so far. *A. fumigatus* encodes one Gwt1 homolog which shares an identity of 38% and similarity of 55.5% with its *S. cerevisiae* homolog. To characterise the role of Gwt1 for growth and viability of *A. fumigatus*, we constructed a conditional mutant by replacing the endogenous promoter of *gwt1* with doxycycline-inducible Tet-On promoter (Helmschrott et al., 2013; Neubauer et al., 2015). This conditional *gwt1_tetOn_* mutant showed significantly reduced radial growth under repressive conditions, while addition of doxycycline fully restored wt-like growth (Fig. 1 A).

**Figure 1.**
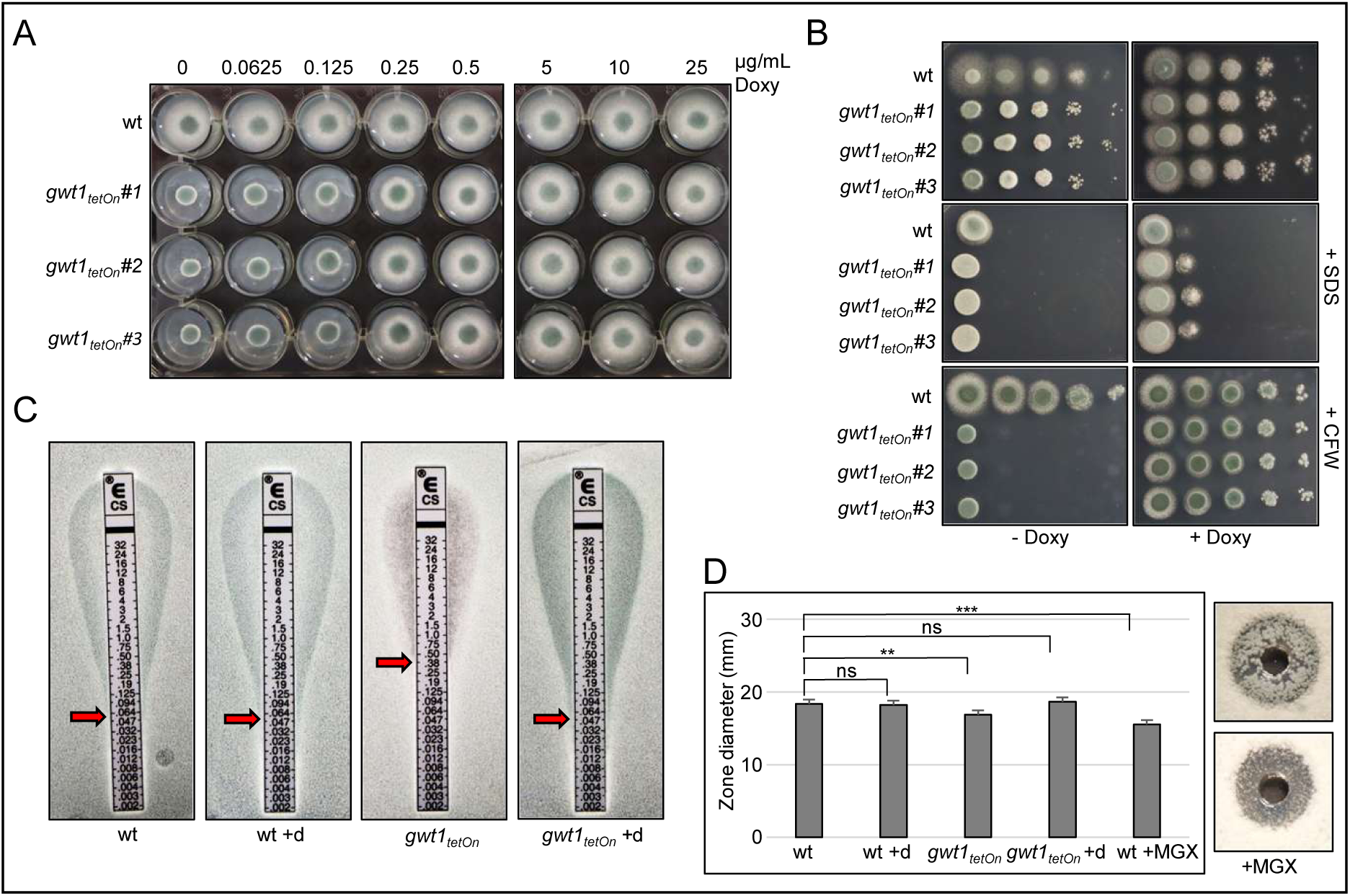
Phenotypic characterisation of a conditional *A. fumigatus gwt1* mutant. (A) 1.5 × 10^3^ conidia of wild-type (wt) or three independent conditional *gwt1_tetOn_* mutants (#1-3) were spotted onto Sabouraud agar (SAB) supplemented with the indicated concentrations of doxycycline (Doxy) and incubated for 28 h at 37 °C. (B) In a series of 10-fold dilutions derived from a starting suspension of 5 × 10^7^ conidia per ml of the indicated strains, aliquots of 3 μl were spotted onto SAB agar plates. Where indicated, media was supplemented with 15 μg ml^-1^ Doxy, sodium dodecyl sulfate (SDS) 0.02% (w/v) or calcofluor white (CFW) 40 μg ml^-1^. Plates were incubated at 37 °C for 30h, except for CFW-supplemented plates which were incubated for 48 h. (C) A total of 1.0 × 10^6^ conidia of the indicated strains were spread on SAB agar plates. When indicated, SAB was supplemented with 15 μg ml^-1^ Doxy (+d). Caspofungin Etest strips were applied, and the agar plates were incubated at 37 °C. Representative photos were taken after 30 hours. (D) 4 × 10^5^ conidia of the indicated strains were spread on SAB agar plates. When indicated, medium was supplemented with 0.05 μg ml^-1^ of manogepix (MGX) or 15 μg ml^-1^ Doxy (+d). Fifty microliters of caspofungin (100 μg ml^-1^) were applied in the punch holes of each agar plate. The inhibition zone diameters of three independent replicates per condition were measured, and images were acquired after 40 h of incubation at 37°C. Statistical significance (*p ≤ 0.05, **p ≤ 0.01, ***p ≤ 0.001) was calculated using the one-way ANOVA test with Dunnett’s post hoc test.

Gwt1 inhibition would be expected to impair the cell wall integrity of fungi. We therefore tested the susceptibility of the mutant to the cell wall perturbing agents calcofluor white, sodium dodecyl sulfate (SDS) and the echinocandin caspofungin. As shown in Fig. 1 B, repression of *gwt1* resulted in a dramatically increased susceptibility to the chitin-binding agent calcofluor white. The susceptibility to the detergent SDS was not increased under repressive conditions. However, under induced conditions, the conditional mutant appeared slightly more resistant (Fig. 1 B). Surprisingly, reduced *gwt1* expression resulted in an increased resistance to caspofungin (Fig. 1 C). Similarly, an increased resistance to caspofungin was observed when Gwt1 was chemically inhibited with manogepix in the wild-type (Fig. 1 C).

### Overexpression of *gwt1* causes manogepix resistance

Overexpression of the drug target can promote resistance to antimicrobial agents. This is well-demonstrated by the most common azole resistance mechanism in *A. fumigatus*, which is a tandem repeat (TR) in the promoter of *cyp51A*, resulting in overexpression of CYP51 (Nywening et al., 2020). In contrast, overexpression of the echinocandin-target β-1,3-glucan glucan synthase is not an established resistance mechanism to echinocandins and rather increases susceptibility to echinocandins (Lee et al., 2023; Loiko and Wagener, 2017). Treatment of *A. fumigatus* wild-type with manogepix resulted in a marked change of the morphology of hyphae (Fig. 2), reminiscent of the growth of *Aspergillus* in the presence of fungistatic concentrations of echinocandins (Dichtl et al., 2015). Interestingly, while repression for *gwt1* resulted in increased susceptibility to manogepix, induction of the conditional promoter significantly increases the resistance to the drug (Fig. 2). This is consistent with similar findings in *Candida albicans* where *gwt1* overexpression also resulted in an increased resistance to Gwt1 inhibitors (Liston et al., 2022; Trzoss et al., 2019; Tsukahara et al., 2003). Our results suggest that overexpression of *gwt1* can be a mechanism of manogepix resistance in *A. fumigatus*.

**Figure 2.**
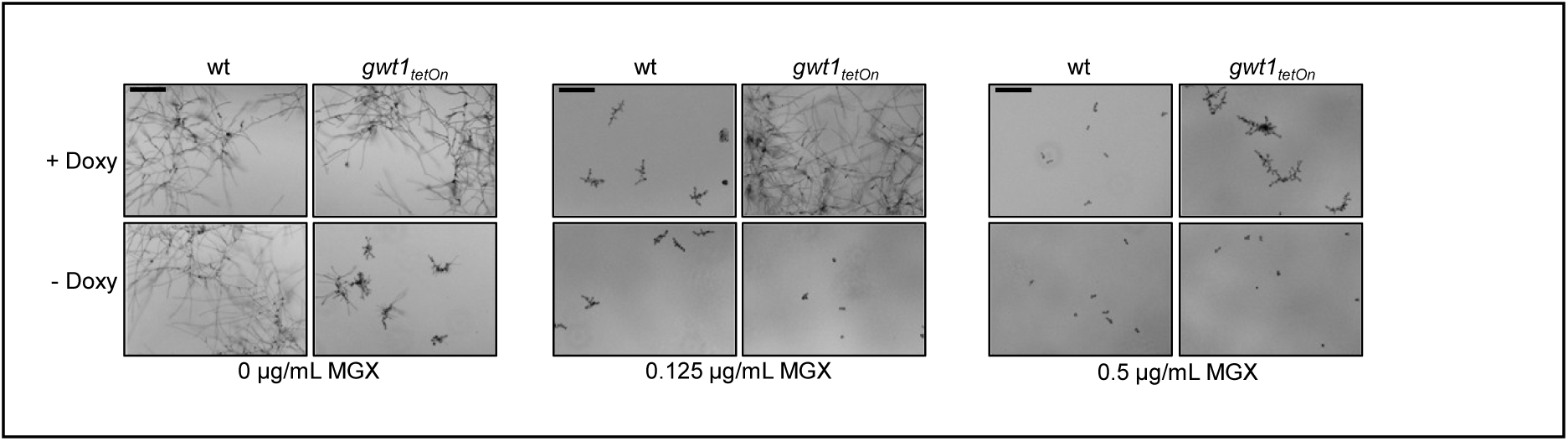
Overexpression of *gwt1* increases resistance to manogepix. Conidia of wild-type (wt) or the conditional *gwt1_tetOn_* mutant were inoculated into wells of a 96-well plate in RPMI*_MOPS_* supplemented with the indicated amount of manogepix (MGX). 15 µg ml^-1^ of doxycycline (Doxy) was supplemented when indicated. Representative images were acquired after 20 h incubation at 37 °C using an automated microscope. Scale bars represent 100 μm and are applicable to all images.

### Manogepix triggers activation of the cell wall salvage response and upregulation of cell wall chitin

Given that GPI-anchored proteins are translocated to the cell membrane to fulfil roles in cell wall remodelling (Ibe and Munro, 2021; Samalova et al., 2020), one may expect that fungi treated with manogepix experience cell wall stress. This would be consistent with the high susceptibility to calcofluor white of the conditional *gwt1_tetOn_* mutant under repressive conditions observed above as well as the occasional cell lysis phenomena observed with the wild type after extensive incubation (Supplementary Fig. 1 A). In fungi, cell wall stress results in activation of a conserved cell wall integrity (CWI) pathway (Dichtl et al., 2016). The promoters of the ɑ-1,3-glucan synthase genes *ags1* in *A. fumigatus* and *agsA* in *Aspergillus niger* are established targets of the canonical CWI pathways in these species (Rocha et al., 2016). To analyse the activation of this pathway by manogepix, a mutant that expresses a reporter construct where firefly luciferase is placed under the control of the *A. niger agsA* promoter was constructed, essentially as described before (Geißel et al., 2018; Ruf et al., 2025). As shown in Fig. 3 A, the cell wall stressor calcofluor white triggered a swift response of the reporter within 20 - 60 minutes. Exposure to the azole antifungal voriconazole resulted in a strong activation of the report after approximately 3 hours (Fig. 3 A), which is in good agreement with our previous findings that activation of the reporter follows the excessive build-up of cell wall (cell wall patches), which is the consequence of the accumulation of the toxic sterol eburicol triggered by CYP51 inhibition (Elsaman et al., 2024; Geißel et al., 2018). The response following manogepix exposure is not as swift as the response following calcofluor white exposure. However, it is much faster than the response following the exposure to voriconazole. In contrast to the voriconazole-induced response, which disappears within the following hours presumably due to the death of the hyphae, the manogepix-induced response steadily increases over time (Fig. 3 A). This demonstrates that manogepix causes cell wall stress and, and is consistent with the low number of cell lysis events (Supplementary Fig. 1 A). This suggests that this antifungal is rather fungistatic and not imminently lethal for *A. fumigatus*.

**Figure 3.**
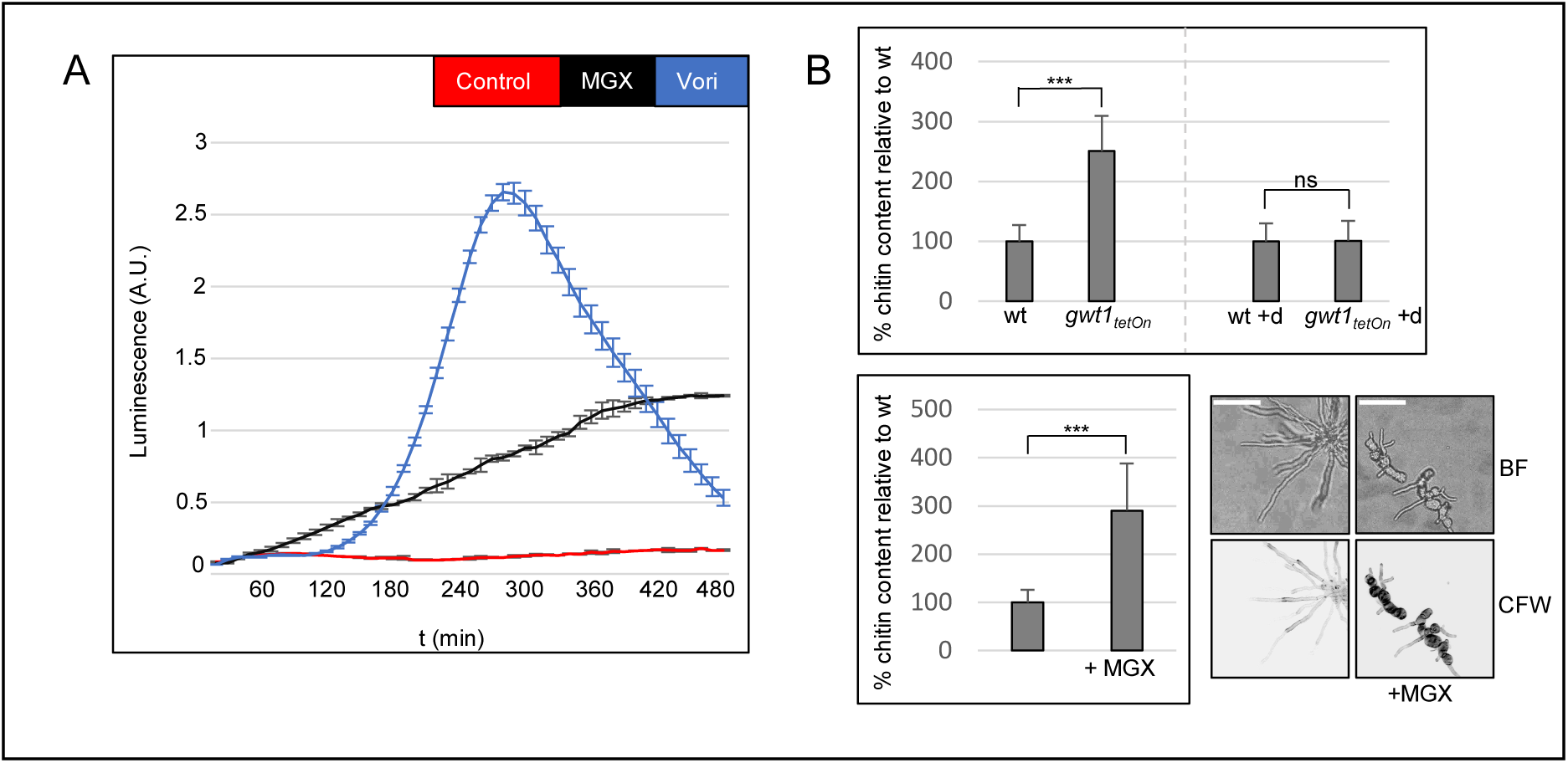
Manogepix-induced stress triggers activation of the cell wall integrity pathway and an increase in cell wall chitin. (A) Conidia of an *A. fumigatus* strain that harbors a luciferase-based cell wall salvage reporter were inoculated in a white 96-well plate in RPMI*_MOPS_* and incubated at 37 °C. After 7 h, medium was supplemented with luciferin and either 0.125 µg ml^-1^ manogepix (MGX), 2 µg ml^-1^ voriconazole (Vori) or no drug (control). Luminescence was monitored in a temperature-controlled microplate reader. The graph depicts the mean luminescence of three technical replicates per condition over time. Error bars represent standard deviations of the three replicates. (B) Wells of an eight-well microscopy slide were co-inoculated with similar numbers of wild-type conidia expressing a cytosolic GFP variant (wt) and conditional *gwt1_tetOn_* mutant conidia (upper bar graph), or inoculated with wild-type conidia without cytosolic GFP (lower bar graph) in RPMI*_MOPS_*. When indicated, medium was supplemented with 15 µg ml^-1^ of doxycycline (+d; upper bar graph) or 0.125 μg ml^-1^ MGX (lower bar graph). Samples were then incubated at 37 °C for 16 h if MGX was added, or 9.5 h if no MGX was added. Hyphae were then fixed and stained with calcofluor white. Samples were subsequently washed three times with sterile water. Chitin levels (calcofluor white fluorescence signal intensity) was then analysed using a confocal laser scanning microscope. At least 10 cell wall cross sections were measured for each condition. Statistical significance (*p ≤ 0.05, **p ≤ 0.01, ***p ≤ 0.001) was calculated with a two-tailed unpaired Student *t* test. The error bars indicate standard deviations. Depicted are representative bright field images (BF) and z-stack projections of optical stacks of the fluorescence signal (chitin) that cover the entire hyphae in focus (CFW). Scale bars represent 50 µm.

It was previously reported that manogepix treatment may result in upregulation of cell wall chitin (Liston et al., 2022). To assess this possibility, we analysed the chitin levels of the manogepix-treated wild-type and the conditional *gwt1_tetOn_* mutant under repressive conditions using the chitin-specific dye calcofluor white (Loiko and Wagener, 2017). As shown in Fig. 3 B and C, reduced expression of *gwt1* as well as manogepix treatment resulted in *A. fumigatus* hyphae with significantly higher chitin levels in their cell walls.

### Cell wall stress response regulators contribute the antifungal activity of manogepix

Our results demonstrated that manogepix treatment induces activation of the CWI pathways, suggesting that the response is important for survival and growth of *A. fumigatus*. To characterise the importance of the CWI pathway-mediated stress response (Dichtl et al., 2016), we analysed the manogepix susceptibility of various mutants with impaired CWI and CWI signaling: Δ*rho2*, a mutant lacking Rho GTPase Rho2 which is involved in cell wall biosynthesis in response to cell wall stress and important for survival of *A. fumigatus* when challenged by human granulocytes or chemical and physical stress conditions such as calcofluor white, Congo red and heat (Dichtl et al., 2012; Ruf et al., 2025). Δ*midA*, a mutant lacking the cell wall stress sensor MidA that is important for survival under very similar stress conditions (Dichtl et al., 2012; Ruf et al., 2025). Δ*wsc1*, a mutant lacking the cell wall stress sensor Wsc1, the absence of which reduces the minimal effective concentration (MEC) of echinocandins (Dichtl et al., 2012). Δ*rho4*, a mutant lacking Rho GTPase Rho4 which is important for septum formation and for survival under various cell wall stress conditions, including when challenged by human granulocytes or azoles antifungals (Dichtl et al., 2015; Geißel et al., 2018; Ruf et al., 2025). And *rho2*-OE, the complemented Δ*rho2* mutant that contains multiple copies of the *rho2* gene due to multiple integration of the complementation plasmid and is more resistant to cell wall stress (Ruf et al., 2025). Susceptibility testing of the wild-type and the CWI mutants was performed following European Committee on Antimicrobial Susceptibility Testing (EUCAST) antifungal susceptibility testing guidelines, with several modifications to allow for a more detailed evaluation of the of the impact on growth and morphology, such as microscop ic imaging after 40 h and using a lower inoculum of spores (conidia), or adding the metabolic indicator resazurin, to estimate the biomass. The MEC of manogepix, the concentration at which a significant morphological switch occurred and hyphal clusters formed, was 0.125 µg ml^-1^ for wild-type *A. fumigatus* (Fig. 4 A). Surprisingly, a mutant lacking Rho2 seemed to be significantly more tolerant to manogepix, showing increased hyphal growth above the MEC compared to wild-type both microscopically and based on resazurin metabolism (Fig. 4 A-C and Supplementary Fig. 2). Notably, distinct morphological differences between the wild-type and Δ*rho2* hyphae were observed when exposed to manogepix, with the Δ*rho2* mutant showing significantly longer and more expanded hyphae, an effect that varied slightly in strength depending on the culture media (Fig. 4 A and C). This is in marked contrast to growth of the Δ*rho2* mutant in the presence of caspofungin, where the opposite effect was observed (Supplementary Fig. 3). Similarly to the Δ*rho2* mutant, a mutant lacking MidA appeared to form slightly longer and more expanded hyphae than to wild-type in the presence of manogepix, although this microscopic finding was not reflected in enhanced overall resazurin metabolism (Fig. 4 A-C and Supplementary Fig. 2). Again, the Δ*midA* mutant appeared to grow slightly worse than the wild-type in the presence of caspofungin (Supplementary Fig. 3). While the Δ*rho2* and Δ*midA* mutants were less constrained in growth in the presence of manogepix, they tended to form more rounded swellings at an earlier stage compared to wild-type at high manogepix concentrations after prolonged incubation, which were prone to cell lysis (Fig. 4 D). Interestingly, osmotically stabilizing the medium with sorbitol partially rescued this growth impairment of the Δ*rho2* mutant at high manogepix concentrations (Supplementary Fig. 1 B).

**Figure 4.**
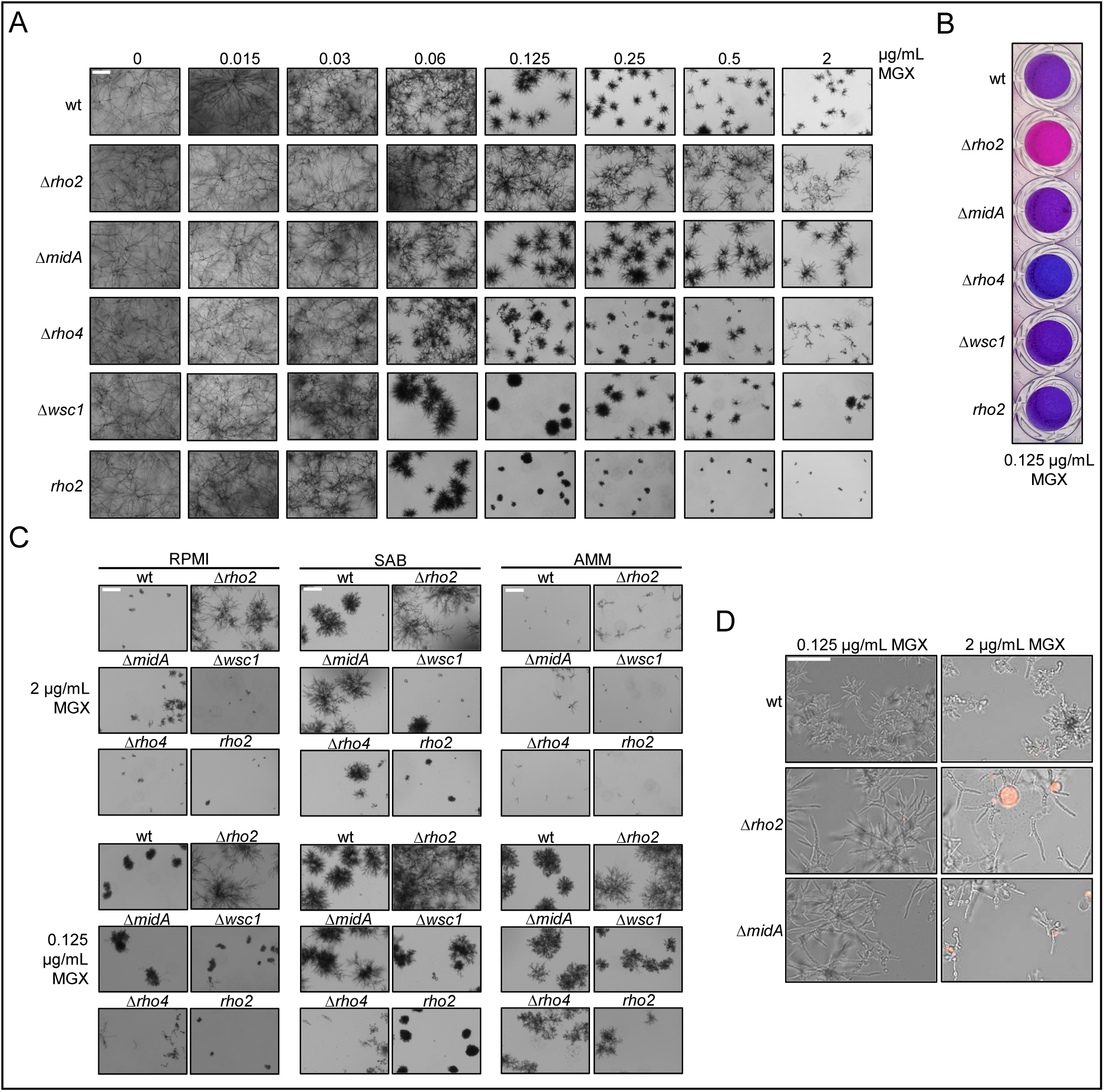
Cell wall stress response regulators Rho2 and MidA enhance the antifungal efficacy of manogepix. (A and B) Conidia of wild-type (wt) and the indicated mutants were inoculated in RPMI*_MOPS_* into wells of a 96-well plate (1 × 10^3^ conidia per well). Medium was supplemented with the indicated concentrations of manogepix (MGX). After 72 h incubation at 37 °C, representative images were acquired using an automated microscope (A) or medium was additionally supplemented with 0.002% (w/v) resazurin, and representative photos taken after a further 16 h incubation at 37 °C (B). (C) Conidia of the indicated strains were inoculated into wells of a 96 well plate, in either RPMI*_MOPS_* (RPMI), Sabouraud (SAB), or *Aspergillus* minimal media (AMM). Medium was additionally supplemented with either 0.125 or 2 µg ml^-1^ of MGX. Representative images were acquired after 48 h incubated at 37 °C using an automated microscope. (D) Conidia of the indicated strains were inoculated in RPMI*_MOPS_* supplemented with 1 µg ml^-1^ propidium iodide and the indicated concentrations of MGX. After 20 h incubation at 37 °C, representative images were acquired using an automated microscope. Depicted are representative images of overlays of the RFP and brightfield channels. (A - C) Scale bars represent 100 µm and are applicable to all images in the respective panels.

Overexpression of *rho2* coincided with an increase in susceptibility to manogepix (1-fold reduced MEC), less tolerance and much smaller, more compact colonies, and reduced resazurin metabolism compared to wild-type (Fig. 4 A-C and Supplementary Fig. 2). As shown in Fig. 4 A and Supplementary Fig. 2, manogepix significantly impaired the growth but also the survival of the Δ*rho4* mutant compared to wild-type, suggesting that manogepix causes a partial fungicidal effect, similarly as it was previously observed for the echinocandin caspofungin (Supplementary Fig. 3 and (Dichtl et al., 2015)). The Δ*wsc1* mutant was slightly more susceptible to manogepix (1-fold reduced MEC), and the mycelium of the emerging microcolonies appeared slightly more compact compared to wild-type (Fig. 4 A-C).

Taken together, these results demonstrated that the lack of Rho2 and - to some extent - the lack of MidA improves growth of *A. fumigatus* in the presence of manogepix. In contrast, the lack of Wsc1 or Rho4 as well as the overexpression of Rho2 slightly increase the susceptibility of this pathogen to manogepix.

### The manogepix-induced increase in cell wall chitin depends on Rho2 and MidA

We have recently shown that Rho2 and MidA are both important for *A. fumigatus* hyphae to withstand attacks from human granulocytes and that activation of Rho2 triggers cell wall remodeling and an increase in cell wall chitin (Ruf et al., 2025). Furthermore, we found that overexpression or constitutive activation of Rho2 significantly reduces the growth rate of *A. fumigatus* (Ruf et al., 2025). We therefore speculated that the lack of Rho2 and (possibly) of MidA hampers the ability of *A. fumigatus* to upregulate cell wall chitin in response to manogepix-induced stress. To test this hypothesis, we exposed CWI mutants to manogepix. Chitin levels were subsequently analysed by staining the cell wall chitin with calcofluor white. The fluorescence signal intensity of numerous cell wall cross sections were measured and compared relative to the wt. As shown in Supplementary Fig. 4, the CWI mutants had chitin levels similar to wild-type under normal growth conditions. However, after manogepix exposure, the lack of Rho2 and to a lesser degree the lack of MidA resulted in a significant decrease of cell wall chitin in *Aspergillus* hyphae compared to the wild-type (Fig. 5). In contrast, neither the lack of Wsc1 or Rho4 seemed to influence the cell wall chitin levels in hyphae after manogepix exposure (Fig. 5). Compared to wild-type hyphae exposed to manogepix, Rho2-overexpressing hyphae showed comparable chitin levels; a minor observed increase was not statistically significant (Fig. 5).

**Figure 5.**
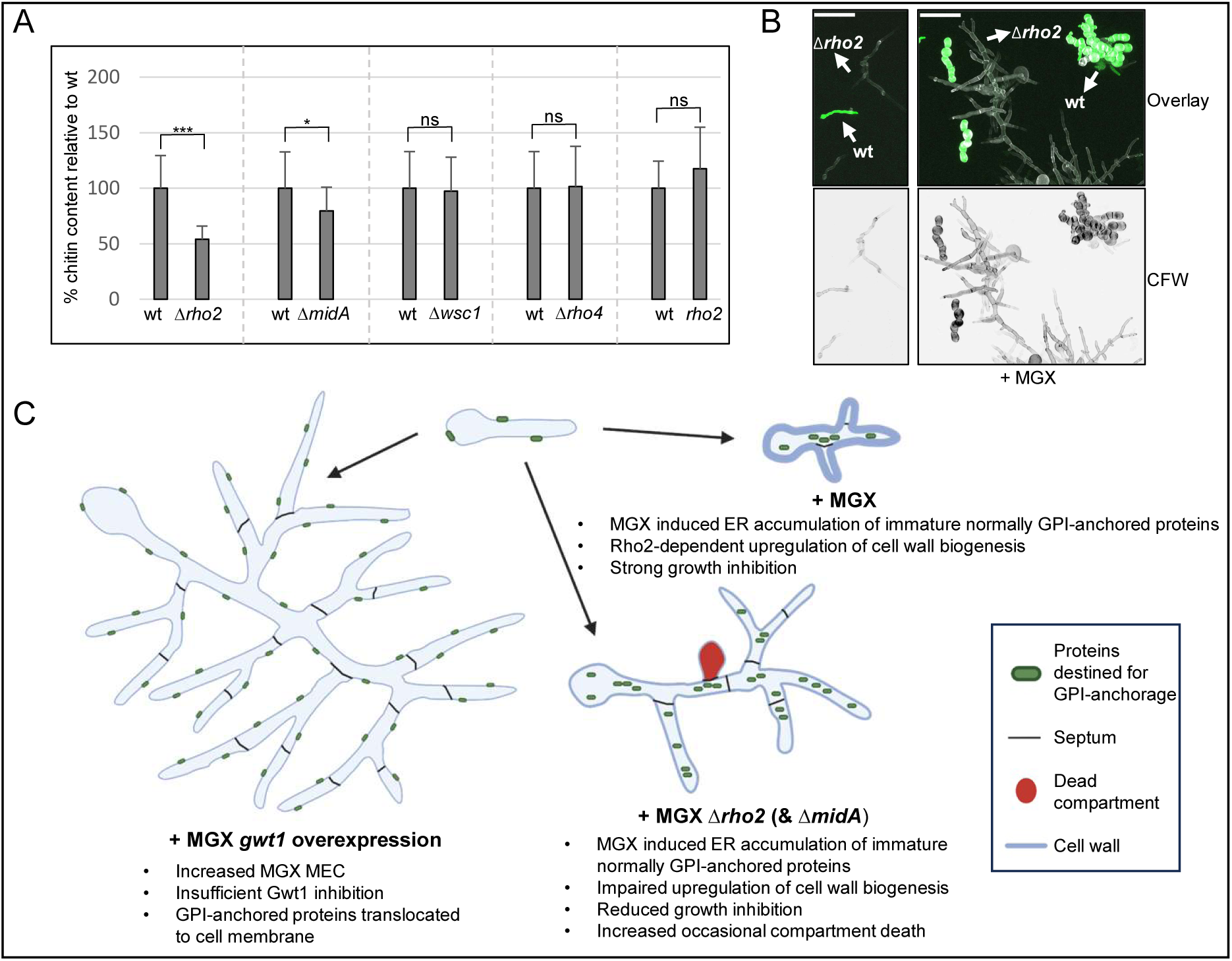
Rho2 and MidA regulate the manogepix-induced increase in cell wall chitin. (A and B) Wells of an eight-well microscopy slide were co-inoculated with similar numbers of wild-type conidia expressing a cytosolic GFP variant (wt) and conidia of the Δ*rho2*, Δ*midA*, Δ*wsc1*, Δ*rho4*, or *rho2* mutant strains in RPMI*_MOPS_* supplemented with 0.125 μg ml^-1^ manogepix (MGX). After 9.5h (B; no MGX) or 16 h (A and B; with MGX) incubation at 37 °C, hyphae were fixed and stained calcofluor white. (A) Chitin levels (calcofluor white fluorescence signal intensity) was then analysed using a confocal laser scanning microscope. At least 10 cell wall cross sections were measured for each condition. Statistical significance (*p ≤ 0.05, **p ≤ 0.01, ***p ≤ 0.001) was calculated with a two-tailed unpaired Student *t* test. The error bars indicate standard deviations. (B) Depicted are representative z-stack projections of optical stacks of the calcofluor white fluorescence signal (chitin; CFW) and overlays of these together with z-stack projections of optical stacks of the GFP fluorescence signal that cover the entire hyphae in focus (CFW). Scale bars represent 50 μm and are applicable to all images. (C) Proposed mechanisms that could promote resistance to MGX in *A. fumigatus*. In a wild-type, MGX inhibits Gwt1 which disrupts the biosynthesis of GPI-anchored proteins and results in accumulation of immature precursors in the ER. The lack of GPI-anchored proteins in the cell wall causes cell wall stress, which triggers the activation of MidA and Rho2, which in turn upregulates cell wall biogenesis and leads to additional growth inhibition. If Rho2 or MidA are lacking (Δ*rho2*, Δ*midA*), MGX-induced cell wall stress fails to trigger the Rho2-dependent upregulation of cell wall biogenesis (e.g., chitin), resulting in improved growth compared to wild-type. The impaired cell wall stress response and MGX-induced cell wall damage occasionally cause the lysis of cell compartments. Hyphae septa, the formation of which is dependent on Rho4, mitigate the impact on the mycelium. Overexpression of *gwt1*, leads to an increased abundance of Gwt1 and insufficient Gwt1 inhibition. GPI-anchored proteins successfully translocate to the cell membrane. This results in an increase of the minimal effective concentration (MEC) of MGX. Created in BioRender. Brazil, S. (2025) https://BioRender.com/s8o7rfn.

## DISCUSSION

Fosmanogepix and its active moiety, manogepix, have demonstrated promising activity against many clinically relevant fungal pathogens. Here, we characterized its activity against *A. fumigatus*, which, alongside *Candida* species, is the most relevant cause of life-threatening invasive fungal infections in Western industrialized countries. Both manogepix treatment as well as reduced expression of the *gwt1* gene encoding the manogepix target showed a strong fungistatic activity against *A. fumigatus*. In contrast, inducing expression of *gwt1* in the *gwt1_tetOn_* mutant resulted in increased resistance to manogepix. This is similar to what was previously observed in *C. albicans* and resembles the CYP51 overexpression-mediated resistance mechanism commonly found in azole-resistant *A. fumigatus* (Liston et al., 2022; Nywening et al., 2020; Trzoss et al., 2019; Tsukahara et al., 2003).

In agreement with previous observations (Miyazaki et al., 2011), manogepix treatment and reduced expression of *gwt1* caused an aberrant growth morphology of the emerging fungal colonies, which was reminiscent of the effects induced by echinocandins. Echinocandins inhibit the biosynthesis of the fungal cell wall polymer β-1,3-glucan which leads to a weakened cell wall. While *A. fumigatus* is able to survive without β-1,3-glucan, frequent cell lysis phenomena of germinating conidia and compartments of hyphae can be observed (Dichtl et al., 2015; Loiko and Wagener, 2017; Moreno-Velásquez et al., 2017). Since some GPI-anchored proteins are important for cell wall biogenesis, manogepix treatment may also lead to an impaired cell wall. In agreement with this, both our study and the literature provide ample evidence that inhibition of Gwt1 function leads to significant cell wall stress and cell wall alterations: 1) Reduced expression of *gwt1* as well as manogepix treatment led to build-up of cell wall chitin, similar as it was observed recently in manogepix-treated *C. albicans* (Liston et al., 2022), a cellular response typically seen under cell wall stress (Ibe and Munro, 2021). 2) Reduced expression of *gwt1* resulted in increased susceptibility to the cell wall-perturbing agent calcofluor, which is in agreement with the increased calcofluor white susceptibility of a *S. cerevisiae* strain with impaired Gwt1 function (temperature sensitive *gwt1-20* mutant) (Umemura et al., 2003). Finally, 3) manogepix exposure triggered a constant activation of the luciferase-based cell wall stress reporter. Based on these results, we investigated how the cell wall integrity and its determinants affects growth and survival of *A. fumigatus* when exposed to manogepix.

Interestingly, the manogepix-induced cell wall alterations and cell wall stress alone did not translate into many cell lysis phenomena in the wild-type. While we could observe occasional death of fungal compartments, similar as recently shown by others (Liston et al., 2022), these events were very rare compared to the frequent death of fungal compartments that is commonly observed after treatment with echinocandins (Dichtl et al., 2015; Moreno-Velásquez et al., 2017). Accordingly, the lack of Rho4, which is important for septum formation, did not have such a detrimental impact on survival of the mutant in the presence of manogepix as it has in the presence of echinocandins (Dichtl et al., 2015, 2012; Thorn et al., 2024). Surprisingly, our results demonstrated that a lack of Rho2 or MidA did not result in increased susceptibility to manogepix. In contrast, we observed significantly increased growth in the presence of manogepix when compared to wild-type. In line with this, overexpression of Rho2 had the opposite effect and resulted in reduced growth in the presence of manogepix when compared to wild-type. Very recently, we have shown that Rho2 upregulates cell wall biosynthesis in response to cell wall stress (Ruf et al., 2025). This is essential for *A. fumigatus* to survive cell wall stress, for example when exposed to calcofluor white or when attacked by granulocytes (Ruf et al., 2025). Furthermore, we have shown that overexpression of Rho2 (using a doxycycline-inducible promoter) reduces the growth rates of *A. fumigatus* and that constitutive activation of Rho2 triggers a full growth arrest accompanied by inappropriate excessive formation of cell wall, including chitin (Ruf et al., 2025).

Taken together, these results suggest that manogepix/Gwt1 inhibition leads to a strong activation of the cell wall stress response of *A. fumigatus*, but the actual accompanying cell wall damage does not pose a significant risk for the cell integrity of the mold. However, the cell wall stress response itself leads to an enhanced fungistatic effect through the activation of Rho2, which enhances the overall antifungal activity. The upregulation of the cell wall stress response induced by manogepix could also explain the slight antagonism we observed with caspofungin: an increase of cell wall chitin is known to improve survival of fungi exposed to echinocandins (Ibe and Munro, 2021; Lee et al., 2023; Wagener and Loiko, 2017).

Could mutations that alter or even disrupt the Rho2-dependent cell wall stress response constitute a clinically relevant resistance mechanism against manogepix? Possible, but on the other hand, conditions in the host are probably more harsh than *in vitro* growth conditions. We have previously shown that a functioning cell wall stress response, including Rho2, is important for the virulence of *A. fumigatus* (Dirr et al., 2010; Ruf et al., 2025). Furthermore, a functioning cell wall stress response is important to withstand other antifungals, e.g., azoles and echinocandins (Dichtl et al., 2012; Dirr et al., 2010; this study). Thus, the benefit of increased resistance to manogepix would probably come at the cost of decreased virulence and decreased resistance to other antifungals. Other mechanisms, such as the increased expression of *gwt1* as shown in a previous study (Liston et al., 2022) and above, might therefore represent a more promising strategy for overcoming manogepix treatment.

In summary, based on our findings, we propose the following mechanisms that can undermine the antifungal activity of manogepix against *A. fumigatus in vitro*, and potentially *in vivo* (Fig. 5 C). 1) Overexpression of *gwt1*, where an increased abundance of the manogepix target Gwt1 diminishes the inhibitory activity of manogepix, very similar to the common CYP51 overexpression-based azole resistance mechanism (Nywening et al., 2020). 2) Alterations of the manogepix-triggered cell wall stress response, thereby preventing the Rho2-dependent upregulation of cell wall biogenesis and growth inhibition. These mechanisms should be considered as possible causes of resistance, which may occur naturally or under treatment and could be observed in the future.

## MATERIAL AND METHODS

### Strains, culture conditions and chemicals

A non-homologous end joining-deficient derivative of the *A. fumigatus* strain D141 (AfS35) served as wild-type in this study (Krappmann et al., 2006; Wagener et al., 2008). All strains used in this study are listed in Table 1. The conditional *gwt1* mutant (*gwt1_tetOn_*) was constructed by inserting a doxycycline-inducible promoter system (pkiA-tetOn; pYZ002) before the coding sequence of *gwt1*, essentially as described before (Helmschrott et al., 2013; Neubauer et al., 2015). The cell wall salvage reporter strain was constructed by transforming wild-type with pBG005-phleo as described before (Geißel et al., 2018). Experiments were performed in Sabouraud medium [2% (w/v) agar, 4% (w/v) d-glucose, 1% (w/v) peptone (#LP0034; Thermo Fisher Scientific; Rockford, IL, US), pH 7.0], in RPMI 1640 (with L-glutamine, but without bicarbonate and without phenol red; R8755; Sigma-Aldrich) supplemented with glucose to a final concentration of 2 % (w/v) and with 3-(N-morpholino) propanesulfonic acid (MOPS) at a final concentration of 0.165 mol/L, pH 7.0 (RPMI*_MOPS_*; medium based on the European Committee on Antimicrobial Susceptibility Testing (EUCAST) antifungal susceptibility testing guidelines (Guinea et al., 2022), but without phenol red) or in *Aspergillus* minimal medium (AMM; (Hill and Kafer, 2001)). Solid media were supplemented with 2 % (w/v) agar (214030; BD Bioscience, Heidelberg, Germany). Doxycycline was obtained from Clontech (631311; Mountain View, CA, USA). Resazurin (#R7017), calcofluor white (#F3543), propidium iodide (#P4170), SDS (#L37671), sorbitol (#1.03557), phosphate-buffered saline (#806552) and caspofungin diacetate (#SML0425) were obtained from Sigma-Aldrich (St. Louis, MO, USA). Voriconazole was purchased from Apexbt Technology LLC (#A4320; Houston, TX, USA). Etest strips were purchased from bioMérieux (Marcyl’Etoile, France). Manogepix (#HY18233) was purchased from MedChemExpress (Monmouth Junction, New Jersey, USA). Luciferin was purchased from Promega (E1601; Fitchburg, WI, USA). Stock solutions for caspofungin diacetate (1 mg ml^-1^), manogepix (5 mg ml^-1^) and voriconazole (10 mg ml^-1^) were prepared in dimethyl sulfoxide (DMSO). Working stocks were freshly prepared by dilution in sterile water. Equivalent volumes of DMSO were added to control samples when appropriate.

**Table 1.**
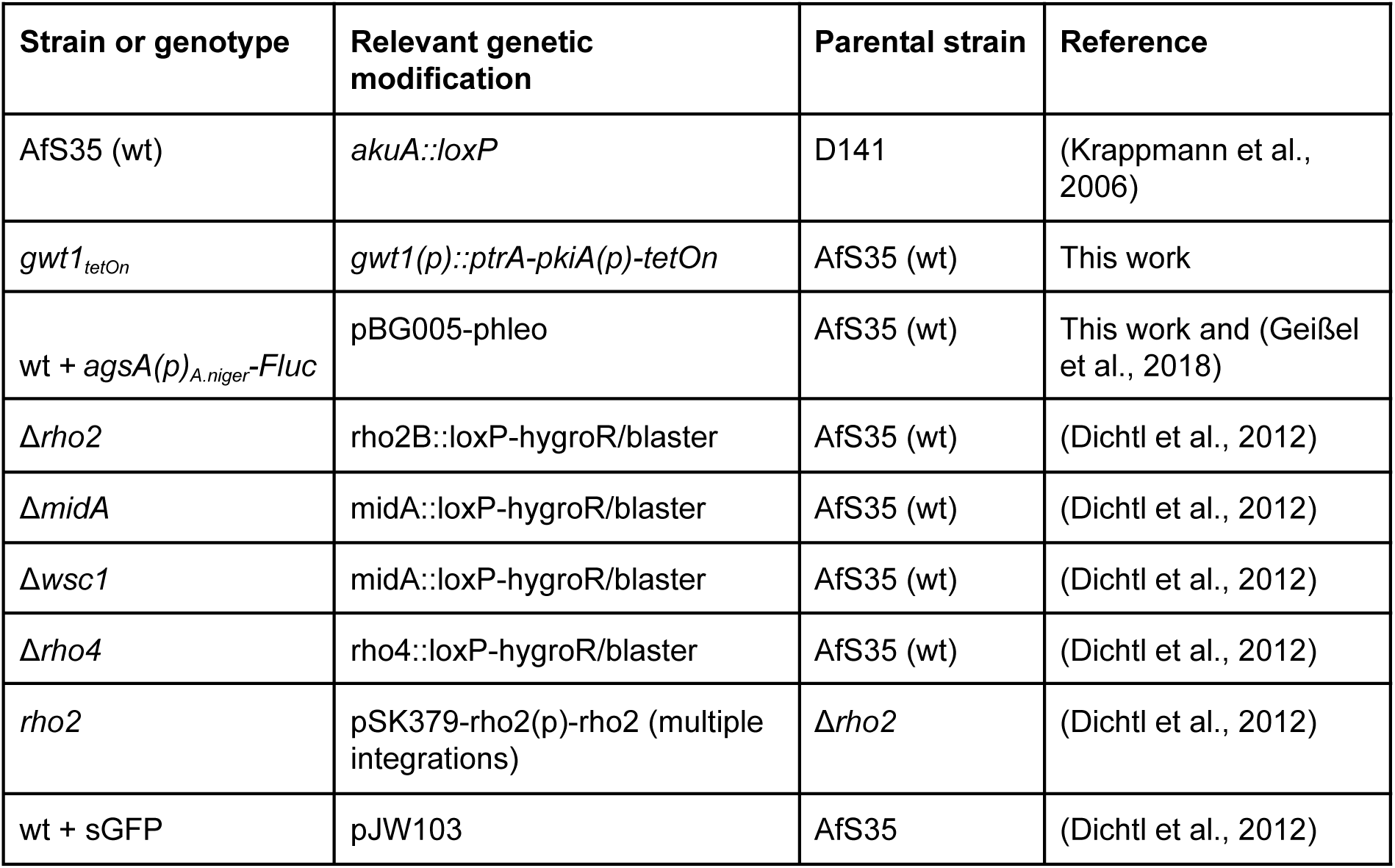
*A. fumigatus* strains relevant for or used in this work.

### Microscopy

The chitin content of cell wall cross sections was analyzed with a confocal laser scanning microscopy (Leica SP8 microscope; Leica Microsystems, Mannheim, Germany). Conidia were inoculated in Ibidi 15 μ-Slide eight-well slides (#80826; Martinsried, Germany). When indicated, samples were fixed with 3.7% (v/v) formaldehyde in phosphate-buffered saline for 3 min. Samples were washed three times with sterile water followed by staining of cell wall chitin with calcofluor white (3.33 µg ml^-1^) for approximately 1 min. Samples were then washed three times with sterile water. The chitin content of cell wall cross sections was analysed using the DAPI channel (excitation 405 nm, emission 410 - 480 nm). An internal wild-type control expressing cytosolic green fluorescent protein (GFP) variant (wt + sGFP) was inoculated in the same well as the *gwt1_tetOn_* mutant, each of the CWI mutants and the Δ*rho4* mutant. This internal wt + sGFP control enabled the comparison of chitin content between wild-type and mutant strains in the same well. Data was anonymised before analysing the chitin content of the cell wall cross sections. Additionally, chitin content of the cross sections in image was analysed prior to unmasking the internal wt + sGFP controls using the GFP channel (excitation 488 nm, emission 502 - 591 nm). At least 10 cross sections were analysed per condition. Statistical significance was calculated using two-tailed unpaired students t-tests.

All other microscopy images were acquired using a Lionheart FX automated microscope (Agilent BioTek; Santa Clara, CA, USA). Conidia were inoculated in clear, flat-bottom, 96-well plates (#3300) purchased from Corning Inc. (NY, USA). When indicated, dead hyphal compartments were stained using 1 µg ml^-1^ propidium iodide and imaged using the RFP channel (excitation wavelength 523 nm; filter cube properties: excitation 531/40 nm, emission 593/40 nm, dichroic mirror 568 nm).

### Cell wall stress reporter assay

To analyse activation of the CWI pathway by manogepix, approximately 2 × 10^4^ conidia per well of the luciferase-based cell wall stress reporter strain wt + *agsA(p)_A.niger_-Fluc* were inoculated in RPMI*_MOPS_*in white 96-well polystyrene microplates (#3917) purchased from Corning Inc. (Corning, NY, USA), essentially as described before (Ruf et al., 2025). After 7 h incubation at 37 °C, medium was supplemented with 0.5 mM luciferin and voriconazole, manogepix or no additional drug. Luminescence was measured over time at 37 °C with a Glomax discover microplate reader (#GM3000; Promega).

### Bioinformatics and calculation of statistical significance

To calculate the percentage identity and similarity between *A. fumigatus* Gwt1 (Afu1g14870) and *S. cerevisiae* Gwt1 (YJL091C) the EMBOSS water Pairwise Sequence Alignment tool was used (Madeira et al., 2024). Statistical significance was calculated using RStudio version 2025.05.0 (Posit Software, Boston, MA, USA).

## AUTHOR CONTRIBUTIONS

J.W. conceived the study. S.B. and J.W. wrote the manuscript draft. I.K., S.Y. and M.C. reviewed and edited the manuscript. S.B., I.K., M.C., and J.W. planned the experiments. S.B., I.K., and S.Y. performed the experiments. Results were analyzed by S.B., I.K. and J.W. All authors read and authorized the manuscript.

## ACKNOWLEDGMENTS

This work was in part supported by Pfizer Healthcare Ireland (Research Grant 73482199), the E.C. Smith Scholarship in Pathology, Trinity College Ireland, and the ERASMUS+ programme.

## DECLARATION OF THE USE OF GENERATIVE ARTIFICIAL INTELLIGENCE

During the preparation of this work the authors used DeepL (deepl.com; Cologne, Germany) for language optimization, i.e., to identify synonyms, nuance phrases, and to correct errors in spelling, grammar, and punctuation.

## FIGURE LEGENDS

**Supplementary Figure 1. Manogepix causes swelling of compartments and death, particularly in the Δ*rho2* mutants, which can be reduced by external osmotic pressure.** (A) Conidia of wild-type were inoculated in RPMI*_MOPS_* supplemented with 1 µg ml^-1^ propidium iodide and 2 µg ml^-1^ of manogepix (MGX). Conidia and newly forming hyphae were imaged over time using an automated microscope. Depicted are representative images of overlays of the RFP and brightfield channels, at the indicated time points. (B) Conidia of wild-type and the Δ*rho2* mutant were inoculated in RPMI*_MOPS_* supplemented with 2 µg ml^-1^ of MGX and 0.5 M sorbitol as indicated. Representative photos were acquired after 72 h incubation at 37 °C. Depicted are representative images of overlays of the RFP and brightfield channels. (A and B) Scale bars represent 50 µm and are applicable to all images.

**Supplementary Figure 2. Metabolic activity of cell wall integrity pathway mutants when exposed to Manogepix.** Conidia of wild-type (wt) and the indicated mutants were inoculated in RPMI*_MOPS_*into wells of a 96-well plate (5 × 10^3^ conidia per well). Medium was supplemented with 0.002% (w/v) resazurin (A) or or manogepix (MGX) 0.125 µg ml^-1^ and 0.002% (w/v) resazurin (B). The plate without MGX was incubated for 18 h at 37 °C. The plate supplemented with MGX was incubated for 24 h at 37 °C. Following incubation resorufin fluorescence (reduced product of resazurin) was analyzed using a microplate reader (excitation wavelength of 520 nm and emission wavelength of 580 - 640 nm). Statistical significance (*p ≤ 0.05, **p ≤ 0.01, ***p ≤ 0.001) was calculated using the one-way ANOVA test with Dunnett’s post hoc test. The error bars indicate standard deviations.

**Supplementary Figure 3. Susceptibility of cell wall integrity pathway mutants to caspofungin.** Conidia of wild-type (wt) and the indicated mutants were inoculated in RPMI*_MOPS_* into wells of a 96-well plate. Medium was supplemented with the indicated concentrations of caspofungin (Caspo). After 72 h incubation at 37 °C, representative images were acquired using an automated microscope. Scale bars represent 100 µm and are applicable to all images.

**Supplementary Figure 4. Cell wall chitin content of cell wall integrity mutants relative to wild-type.** Wells of an eight-well microscopy slide were co-inoculated with similar numbers of wild-type conidia expressing a cytosolic GFP variant (wt) and conidia of the Δ*rho2*, Δ*midA*, Δ*wsc1*, Δ*rho4*, or *rho2* mutant strains in RPMI*_MOPS_*. After 9.5h incubation at 37 °C, hyphae were fixed and stained calcofluor white. Samples were subsequently washed three times with sterile water. (A) Chitin levels (calcofluor white fluorescence signal intensity) was then analysed using a confocal laser scanning microscope. At least 10 cell wall cross sections were measured for each condition. Statistical significance (*p ≤ 0.05, **p ≤ 0.01, ***p ≤ 0.001) was calculated with a two-tailed unpaired Student *t* test. The error bars indicate standard deviations.

