## Supplementary figures and images for "Rho2-dependent cell wall remodeling boosts the fungistatic activity of manogepix against *Aspergillus fumigatus*"

### Supplementary Figures

#### Supplementary Fig. 1

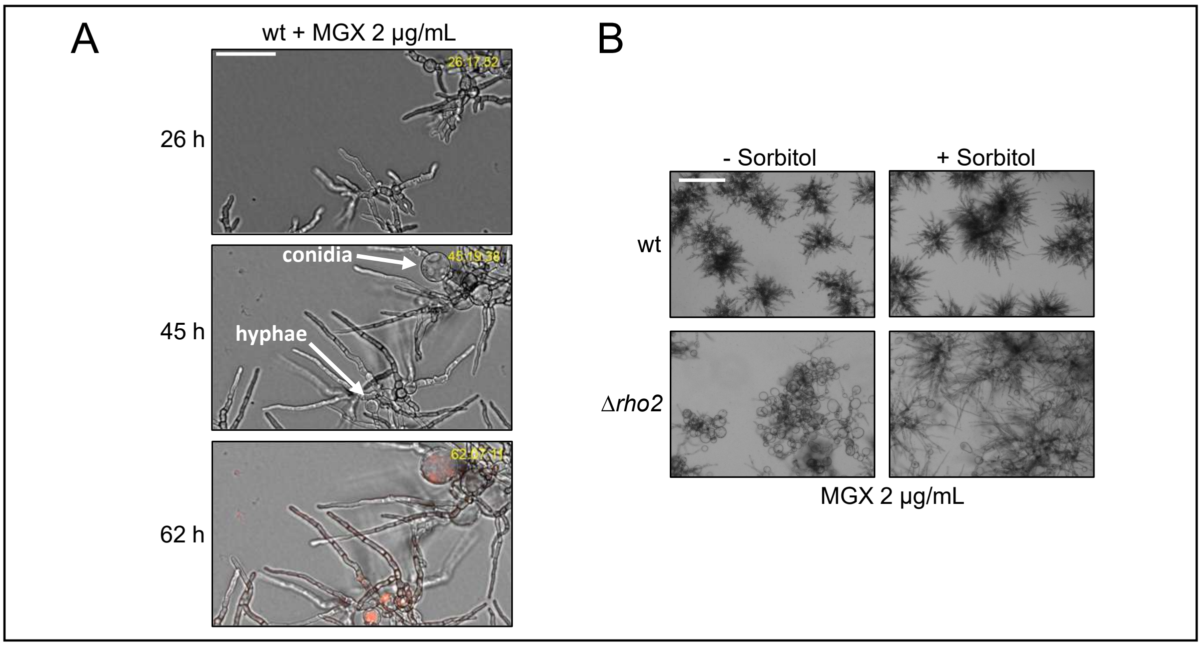

#### Supplementary Fig. 2

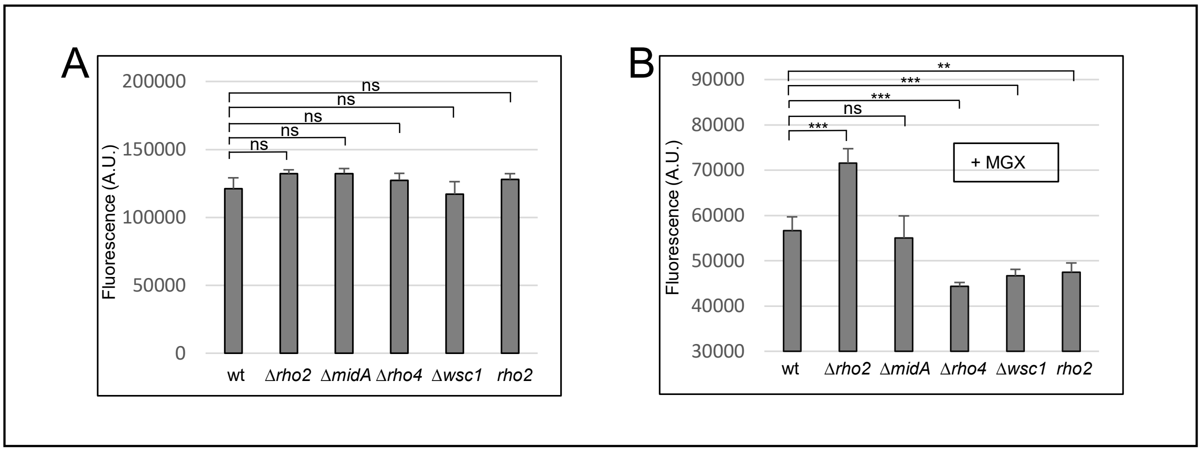

##### Supplementary Fig. 3

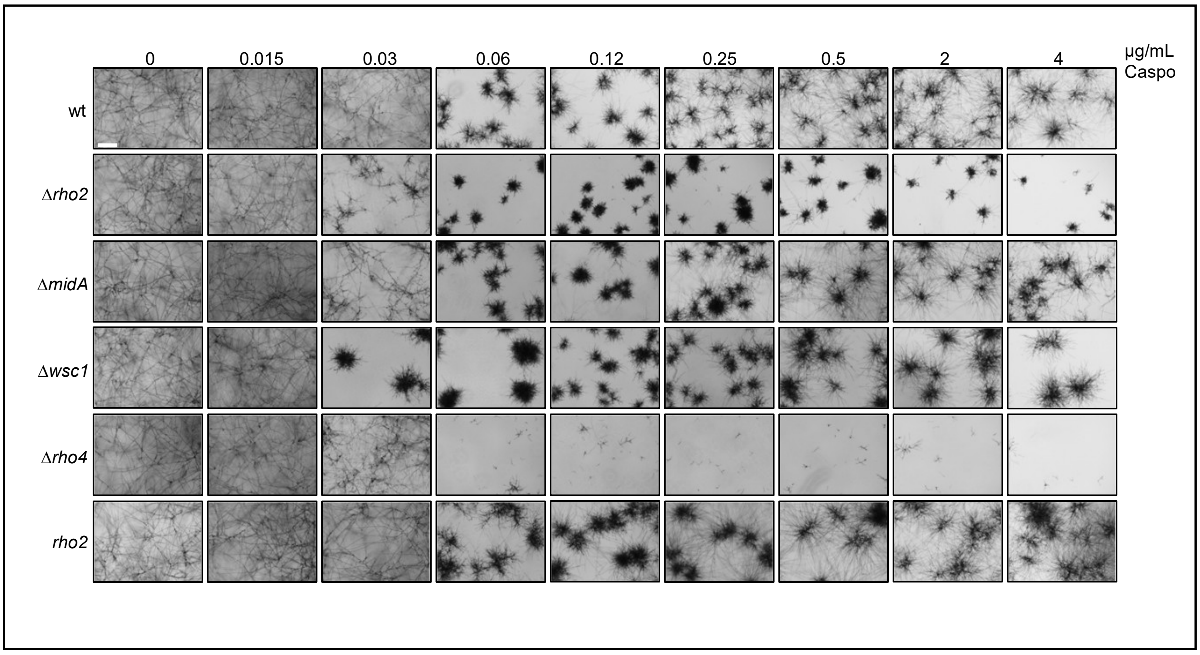

#### Supplementary Fig. 4

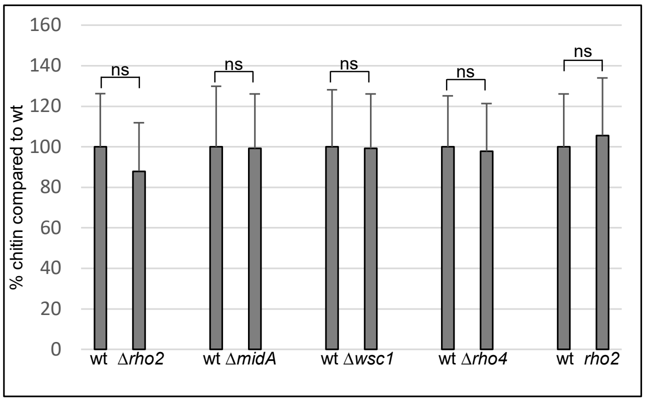
